# Functional connectivity of music-induced analgesia in fibromyalgia

**DOI:** 10.1101/230243

**Authors:** Victor Pando-Naude, Fernando A. Barrios, Sarael Alcauter, Erick H. Pasaye, Lene Vase, Elvira Brattico, Peter Vuust, Eduardo A. Garza-Villarreal

## Abstract

Listening to self-chosen, pleasant and relaxing music reduces pain in fibromyalgia (FM), a chronic central pain condition. However, the neural correlates of this effect are fairly unknown and could be regarded as a more direct measure of analgesia. In our study, we wished to investigate the neural correlates of music-induced analgesia (MIA) in fibromyalgia patients. To do this, we studied 20 FM patients and 20 matched healthy controls (HC) acquiring rs-fMRI with a 3T MRI scanner, and pain data before and after two 5-min auditory conditions: music and noise. We performed resting state functional connectivity (rs-FC) seed-based correlation analyses (SCA) using pain and analgesia-related ROIs to determine the effects before and after the music intervention in FM and HC, and its correlation with pain reports. We found significant differences in baseline rs-FC between FM and HC. Both groups showed changes in rs-FC in several ROIs after the music condition between different areas, that were left lateralized in FM and right lateralized in HC. FM patients reported MIA that was significantly correlated with rs-FC decrease between the angular gyrus, posterior cingulate cortex and precuneus, and rs-FC increase between amygdala and middle frontal gyrus. These areas are related to autobiographical and limbic processes, and auditory attention, suggesting MIA may arise as a consequence of top-down modulation, probably originated by distraction, relaxation, positive emotion, or a combination of these mechanisms.

## INTRODUCTION

Music-induced analgesia (MIA) is defined as the subjective reduction of pain perception after listening to music (Roy et al., 2008), and the effect has been reported in chronic pain conditions such as low back pain, osteoarthritis, and fibromyalgia (Siedliecki et al., 2006; Guétin et al., 2012; Onieva-Zafra et al., 2013). Although the neural correlates of MIA have not been thoroughly studied, endogenous pain inhibition depends on the descending pain modulatory system (DPMS), with areas involved such as: the dorsolateral prefrontal cortex (dlPFC), periaqueductal gray matter (PAG) and rostral ventral medulla (RVM) (Staud, 2012; Tracey et al., 2012). Thus, the possible neural mechanisms of MIA seem to be top-down through the DPMS, secondary to cognitive and emotional mechanisms such as: distraction (Mitchell et al., 2006; Garza-Villarreal et al., 2012), familiarity (Pererira et al., 2011; van der Bosch et al., 2013), emotion (Roy et al., 2012), relaxation and reward (Rhudy et al., 2008; Salimpoor et al., 2013; Hsieh et al., 2014). MIA may be then catalogued as a central type of analgesia, given that the effect seems to originate in the brain and not by peripheral nociceptive receptors (Dobek et al., 2014).

Fibromyalgia (FM) is a chronic pain syndrome of unknown etiology that predominantly affects women, and is characterized by increased sensitivity to somatosensory nociception, and associated with other symptoms such as: sleep disorders, stiffness, fatigue, anxiety, and depression (Wolfe et al., 2010; Napadow et al., 2010; Jensen et al., 2010). There is still no specific treatment for FM and conventional treatment can result in abuse of painkillers, which lead to other co-morbidities (Borchers & Gershwin, 2015). FM patients seem to exhibit a decrease of central inhibition or facilitation of the nociceptive input in the DPMS (Petersel et al., 2010; Brederson et al., 2011; de la Coba et al., 2017), and thus are more sensitive to pain, as well as other types of sensory input such as noise. In consequence, this seems to be reflected by increased function of the pain pathways, increased membrane excitability and synaptic efficacy, as well as reduced neuronal inhibition (Latremoliere & Woolf, 2009).

In FM, several studies have shown morphological and functional characteristics of these patients using different neuroimaging techniques (Sawaddiruk et al., 2017). Specifically, recent resting-state fMRI studies have found alterations in brain connectivity in FM patients (Gracely et al., 2002; Williams & Gracely, 2006; Napadow et al., 2014), involving networks related to pain intensity and analgesia (Napadow et al., 2010; Napadow et al., 2012; Kim et al., 2013; Cummiford et al., 2016). FM patients show increased resting state functional connectivity (rs-FC) of areas related to pain processing, and reduced connectivity in regions involved in pain inhibitory modulation (Cifre et al., 2012). These findings include increased connectivity of insula (INS) and thalamus (THA) (pain-related) with the posterior cingulate cortex (PCC) and medial prefrontal cortex (mPFC) (Default Mode Network-related), as well as decrease connectivity between THA, premotor areas, INS, primary somatosensory cortex (SI), and prefrontal areas (Napadow et al., 2010; Flodin et al., 2014). FM patients have shown increased connectivity of the anterior cingulate cortex (ACC) with INS and basal ganglia; secondary somatosensory cortex (SII) with amygdala (AMYG); and mPFC with PCC. Also, they have shown decreased connectivity of the ACC with AMYG and PAG; THA with INS and PAG; INS with putamen (PUT); PAG with caudate (CAU); SII with primary motor cortex (M1) and PCC; and PCC with superior temporal sulcus (STS) (Cifre et al., 2012; Ichesco et al., 2014; Lazaridou et al., 2017; Truini et al., 2016; Coulombe et al., 2017). Thus, alterations in rs-FC in FM patients appears to involve not only areas related to pain processing (perception and modulation), but also related to somatomotor, executive, limbic, autobiographic, and integration processes.

In terms of MIA, there are no neuroimaging studies in acute or chronic pain, with the exception of our own previous study, where, we showed MIA related to increased BOLD signal amplitude in the angular gyrus (AnG) in FM patients (Garza-ViIlarreaI et al., 2015). Although the changes in amplitude were correlated with the pain self-report, the study did not include healthy controls and functional connectivity changes may be random and unrelated to the analgesic effect. Overall, if the mechanisms behind MIA are related to the DPMS, areas such as the ACC, PAG and INS should show rs-FC changes in FM patients. In this study, we wished to investigate rs-FC patterns of MIA in FM patients, compared to age and sex-matched healthy controls (HC), by means of pain self-report, rs-fMRI and seed-based correlation analyses on the rs-FC of our experimental pain network (e-PNN). We hypothesized that (1) FM patients would show significant differences in rs-FC of areas related to pain processing at rest, and (2) the analgesic effect of music would be associated to changes in rs-FC of areas related to the DPMS in FM patients.

## MATERIALS AND METHODS

### Participants

The study was conducted at the Instituto de Neurobiología of the Universidad Nacional Autónoma de México (UNAM Juriquilla, Queretaro, Mexico). A fibromyalgia group (FM, n = 20, age range = 22 - 70, mean = 46.4, SD = 12.5) and an age-matched control group (HC, n = 20, age range = 21-70, mean = 42.1, SD = 12.5) participated in this study. The inclusion and exclusion criteria for participation in the fMRI experiment are described in **Table 1**. All participants gave their consent verbally and in written form before the experiment. FM patients were asked not to intake painkillers at the day of testing only. The HC group were screened to ensure that they did not experience any type pain at the day of testing. The study was conducted in accordance with the Declaration of Helsinki and approved by Bioethics Committee of the Instituto de Neurobiología, UNAM. Patients received no compensation for participating in the study.

**Table I.** Participant selection criteria.

|  |
| --- |
| **INCLUSION CRITERIA FOR FIBROMYALGIA PATIENTS.** |
| • Meeting the Fibromyalgia 1990 and 2010 criteria (Wolfe et al., 1990, 2010).<br>• Fibromyalgia diagnosed by a trained Rheumatologist.<br>• Spontaneous, continuous and intense pain in daily life (VRS >5 average of a month)<br>• Right-handed. |
| **INCLUSION CRITERIA FOR HEALTHY CONTROLS.** |
| • Healthy Adult.<br>• Right-handed. |
| **EXCLUSION CRITERIA FOR FIBROMYALGIA PATIENTS AND HEALTHY CONTROLS.** |
| • Impossibility to move or walk.<br>• Uncontrolled endocrine problems.<br>• Neurological alterations (i.e. stroke, epilepsy, recent traumatic brain injury).<br>• Auditory problems.<br>• MRI contraindications (i.e., metal prosthetics).<br>• Pregnancy and/or breast-feeding. |
| **ELIMINATION CRITERIA FOR FIBROMYALGIA PATIENTS AND HEALTHY CONTROLS.** |
| • Excessive artifacts in MRI.<br>• Probable pathological findings in MRI. |
| VRS, Verbal Rating Scale. |

### Design and Paradigm

Part of the current data has been previously analyzed and published by Garza-Villarreal et al. (2015), which included FM patients only. In this new study, we included healthy controls and performed a seed-based functional connectivity analyses. Participants answered the Pain Catastrophizing Scale (PCS), the State-Trait Anxiety Inventory (STAI), the Pain Self-Perception Scale (PSP), and the Center for Epidemiologic Studies Depression Scale (CES-D) before the MRI scanning, to establish clinical and behavioral differences between FM patients and HC. To evaluate pain while in the MRI scanner, pain intensity (PI) and pain unpleasantness (PU) were measured only in FM patients **(Figure 1),** using the verbal rating scale (VRS) (0 = no pain, 10 = worst pain possible) (Cork et al., 2004). PI refers to the sensory aspect of pain, whereas PU refers to the evaluative and emotional dimension of pain (Price, 1999). PI and PU were measured immediately before and after each experimental condition. The experimental conditions consisted of five-minute long auditory tracks, either music or pink noise (control), presented while no imaging was acquired, with participants inside the MRI scanner. Prior to the study, participants provided a list of songs or artist that they would like to listen during the experiment. Songs had to be highly pleasant and slow paced. The slow pace was defined as a tempo of < 120 beats per minute (bpm), determined by the researcher using a metronome. Pleasantness was reported by the participant using a 10-point verbal scale (0 = unpleasant, 10 = highly pleasant), and to be selected, the song had to be rated at least 9 - 10 points. When only the artist name was provided, the researcher chose the songs based on two fixed acoustic criteria: consonance (pleasantness), verbally reported by the participant, and slow tempo. Pink noise was selected by a prior pilot study in which several types of noise were presented to healthy participants (Garza-Villarreal et al., 2012). Pink noise resulted as more neutral than other types of noise (i.e. white noise).

Participants listened to the auditory stimuli inside the MRI scanner **(Figure 1),** a period in which no sequences were acquired to minimize unwanted noise. The order of the auditory stimulus presentation was counter-balanced across participants, to avoid any order effect. Auditory stimuli were presented using the NordicNeuroLab AS (Bergen, Norway) MRI-safe headphones. For each session, there were a total of four rs-fMRI acquisitions (rs-fMRI 1, rs-fMRI 2, rs-fMRI 3, rs-fMRI 4) lasting five minutes each, in which participants were instructed to stay alert with eyes opened and a fixation cross was presented on a screen. A wash-out condition was presented between the second and third functional acquisition, and consisted of watching a video documentary with sound (i.e., a biography of Bill Gates), period in which structural imaging was acquired **(Figure 1).** The purpose of the wash-out condition was to avoid analgesic or cognitive cross-over effects. Visual stimuli (fixation and wash-out condition) were presented in a screen projected through a mirror mounted on the MRI head coil, using the software VLC Media Player (http://videolan.org). A total of five conditions were defined for the statistical analysis: baseline (BL), pre-control (Cpre), post-control (Cpos), pre-music (Mpre), and post-music (Mpos). The BL condition was defined as the first rs-fMRI sequence acquired, and it was used to analyze differences in brain functional connectivity between groups (FM and HC) before the music intervention.

**Figure 1.**
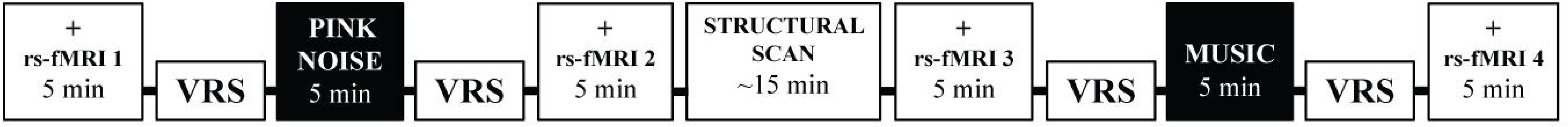
Experimental rs-fMRI Paradigm. Experimental conditions were pink noise and music. Image acquisitions were performed before and after each experimental condition, in which participants were instructed to stay alert with eyes opened and fixated on a white cross displayed on the center of a black background presented on a screen. Pain Intensity and Pain Unpleasantness was reported by fibromyalgia patients only, before and after each experimental condition. The washout condition was executed during the "Structural Scan" period. PINK NOISE, control condition; VRS, pain verbal rating scale; rs-fMRI, resting state functional magnetic resonance imaging; +, eye fixation.

### Procedure

FM patients were recruited through a Fibromyalgia support group and from the General Hospital of the Health Government Department (Hospital General de la Secretaría de Salud) both located in Queretaro, Mexico. HC were recruited using flyers placed in the Instituto de Neurobiología, and with the help of students and workers of the same institute. Potential participants were informed and interviewed by phone to make sure they met the inclusion criteria. After participants were confirmed to be eligible and accepted to participate in the study, they were asked for songs they would like to listen to during the study, that would fit the characteristics described in the previous section. Before the MRI scans, they were briefed about the study to make sure they understood the procedure and implications. Participants then answered the behavioral questionnaires described above. During the MRI scanning, participants in the FM group rated their spontaneous pain immediately before and after each auditory condition. The HC group experienced no pain, thus, pain was not measured.

### MRI Data Acquisition

The image acquisition was performed with a 3.0 Tesla GE Discovery MR750 scanner (HD, General Electric Healthcare, Waukesha, WI, USA) and a commercial 32-channel head coil array. High-resolution T1-weighted anatomical images were obtained using the FSPGR BRAVO pulse sequence: plane orientation = sagittal, TR = 7.7 ms, TE = 3.2 ms, flip angle = 12°, matrix = 256 × 256, FOV = 256 mm^2^, slice thickness = 1 mm, number of slices = 168, gap = 0 mm, slice order = interleaved, view order = bottom-up. A gradient echo sequence was used to collect rs-fMRI data using the following parameters: plane orientation = axial, TR = 3000 ms, TE = 40 ms, flip angle = 90°, matrix = 128 × 128, FOV = 256 mm^2^, slice thickness = 3 mm, voxel size = 2 × 2 mm, number of slices = 43, gap = 0 mm, slice order = interleaved, view order = bottom-up. The total time for each rs-fMRI sequence was 5 minutes with a total of 100 brain volumes acquired per run, with 4 runs per subject. During the scanning, patients were not given any instructions about the music or pink noise, but were instructed to stay alert, to keep their eyes open without thinking anything in particular. All images were saved in DICOM format, anonymized and converted to NIFTI format using dcm2nii from MRIcron (Rorden and Brett, 2000).

### Statistical Analysis of Questionnaires and Pain Measures

Descriptive and inferential statistics of data and plots were performed using R software (R Core Team, 2013) and the “ggplot2” package of R (Wickham, 2009). To establish behavioral differences between experimental groups (FM and HC), a two-tailed unpaired *t-student* test was performed on the results from pain self-perception, pain catastrophizing, anxiety, and depression questionnaires. PI and PU scores of the FM group were not normally distributed, therefore, non-parametric two-tailed paired analyses were performed with the *Mann-Whitney Rank* test. This analysis was performed in the difference of variables ΔPI (pre-post PI) and ΔPU (pre-post PU) between the two experimental conditions (music and pink noise). Three FM patients were excluded from this analysis due to technical difficulties to record the pain rating data.

### Functional Connectivity Analysis

The rs-fMRI data was preprocessed and analyzed using the CONN Toolbox for Matlab (Functional Connectivity Toolbox, Gabrieli Lab., 2015). Structural and functional images were imported into CONN and the preprocessing pipeline included: realignment, slice-timing correction, structural segmentation and spatial normalization (simultaneous Gray/White/CSF segmentation and normalization to the MNI space), outlier detection (ART-based identification of outlier scans for scrubbing; subject motion correction = 2.5 mm and global signal Z-value threshold = 3), and smoothing (spatial convolution with a Gaussian kernel with FWHM = 5 mm). Nuisance variables were regressed out using the general lineal model. Signal timeseries were band-pass filtered between 0.008 and 0.09 Hz. Nuisance variables included six motion variables, and principal components of white matter and cerebrospinal fluid, a method referred as aCompCor (Muschelli et al., 2014). The aCompCor avoids artefactual anticorrelations introduced by global signal regression and reduces artifact by physiological signals. To evaluate resting state functional connectivity (rs-FC), seed-based correlation analyses (SCA) were performed against whole brain. We defined an experimental Pain Neural Network (e-PNN) pre-hoc using the following brain areas in both hemispheres: ACC, AnG, AMYG, primary auditory cortex (BA41), CAU, globus pallidus (GP), PUT, INS, mPFC, PAG, PCC, M1, SI, SII, supplementary motor area (SMA), STS, and THA **(Figure 2, Supplementary Table I).** The e-PNN ROIs were defined according to several pain studies, both experimental and clinical (Gracely et al., 2002; Zaki et al., 2007; Baliki et al., 2008; Burgmer et al., 2009; Cifre et al., 2012; Garza-Villarreal et al., 2015), as well as a neuroimage atlases (Harvard-Oxford atlas FSLview; Juelich Histological Atlas FSLview), and Neurosynth (Yarkoni et al., 2011) using “pain” and “chronic pain” as search terms for the later. The 34 ROIs were created using fslmaths (FMRIB Software Library v5.0, Analysis Group, FMRIB, Oxford, UK), with a 5-mm kernel sphere each in MNI space. We then performed seed-to-voxel correlation analyses for each seed independently using the CONN Toolbox. For the statistical analysis, first we compared BL condition between experimental groups (FM vs HC) using the following covariates: age, years with FM diagnosis, and anxiety and depression symptoms. Then, we performed a within-subject contrast of the different conditions: Cpre vs Cpos, and Mpre vs Mpos. All analyzed contrasts were performed using t-tests corrected for multiple comparisons using the false discovery rate (FDR) at q = 0.05 for each test and for each cluster. To determine if the analgesic effect of music correlates with rs-FC results in FM patients, the resulting values of ΔPI and ΔPU scores, along with the rs-FC contrast results of Mpre vs Mpos (Δrs-FC) values were transformed to Z-values, and a two-tailed Pearson’s correlation analysis was performed, considering it significant at α = 0.05.

**Figure 2.**
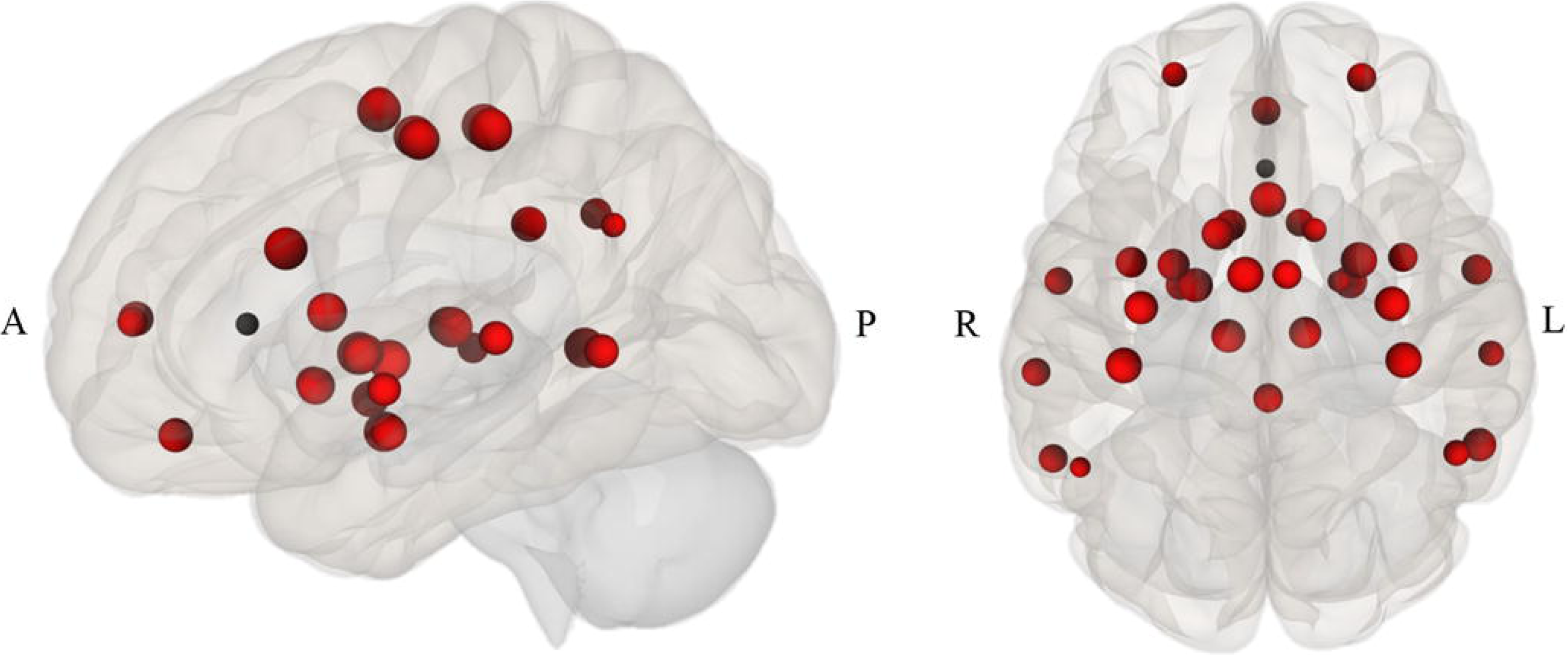
rs-fMRI seed-based correlation analysis. Experimental Pain Neural Network (e-PNN): ACC, anterior cingulate cortex; PCC, posterior cingulate cortex; BA41, primary auditory cortex; AMYG, amygdala; AnG, angular gyrus; CAU, caudate; GP, globus pallidus; PUT, putamen; INS, insular cortex; mPFC, medial prefrontal cortex; PAG, periaqueductal gray matter; M1, primary motor cortex; SI, primary somatosensory cortex; SII, secondary somatosensory cortex; SMA, supplementary motor area; STS, superior temporal sulcus; THA, thalamus. A, anterior; P, posterior; R, right; L, left.

## RESULTS

### Questionnaires and Pain Measures

As expected and in accordance with previous FM behavioral studies, pain self-perception, pain catastrophizing, anxiety and depression were significantly different between FM and HC **(Table II).** Patients with FM present greater pain self-perception (<0.001), pain catastrophizing (p<0.001), anxiety (p<0.001) and depression (p<0.001) symptoms than HC. The FM group reported significantly less pain after listening to music only: ΔPI (W=60, p=0.002) and ΔPU (W=65.5, p=0.004).

**Table II.**
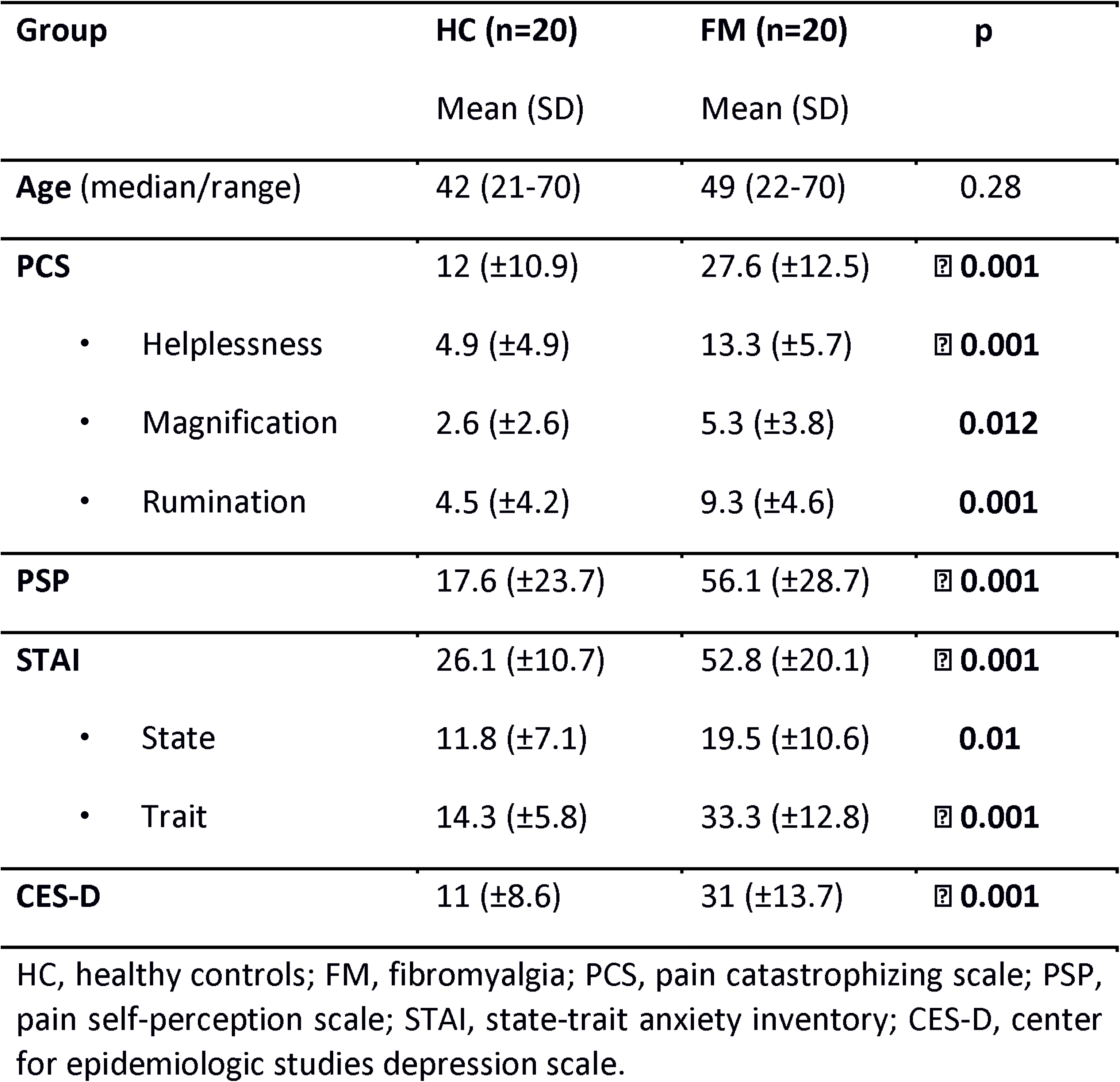
**Descriptive and Inferential Statistics of Behavioral Questionnaires**. HC, healthy controls group; FM, fibromyalgia group; PCS, pain catastrophizing scale; PSP, pain self-perception scale; STAI, state-trait anxiety inventory; CES-D, center for epidemiologic studies depression scale.

| Group | HC (n=20) | FM (n=20) | p |
| --- | --- | --- | --- |
|  | Mean (SD) | Mean (SD) |  |
| Age (median/range) | 42 (21-70) | 49 (22-70) | 0.28 |
| **PCS** | 12 (±10.9) | 27.6 (±12.5) | **0.001** |
| • Helplessness | 4.9 (±4.9) | 13.3 (±5.7) | **0.001** |
| • Magnification | 2.6 (±2.6) | 5.3 (±3.8) | **0.012** |
| • Rumination | 4.5 (±4.2) | 9.3 (±4.6) | **0.001** |
| **PSP** | 17.6 (±23.7) | 56.1 (±28.7) | **0.001** |
| **STAI** | 26.1 (±10.7) | 52.8 (±20.1) | **0.001** |
| • State | 11.8 (±7.1) | 19.5 (±10.6) | **0.01** |
| • Trait | 14.3 (±5.8) | 33.3 (±12.8) | **0.001** |
| **CES-D** | 11 (±8.6) | 31 (±13.7) | **0.001** |

HC, healthy controls; FM, fibromyalgia; PCS, pain catastrophizing scale; PSP, pain self-perception scale; STAI, state-trait anxiety inventory; CES-D, center for epidemiologic studies depression scale.

### Functional Connectivity

#### Baseline Condition

The FM vs HC contrast analysis for the BL condition, revealed that FM patients show higher connectivity of the following seeds: left AnG with precuneus (PCN), left paracingulate gyrus (PaCiG), right temporal pole (TP), left anterior middle temporal gyrus (MTG), subcallosal cortex (SubCalC), left frontal pole (FP), and right CRBL; right GP with left inferior frontal gyrus (IFG); left GP with left supramarginal gyrus (SMG), right SMA, and right superior parietal lobe (SPL); left mPFC with left AnG and right FP; right PCC with right PaCiG, PCC, right FP, and left AnG; left PCC with PCN, left superior lateral occipital cortex (SLOC), right SLOC, and right PaCiG; left THA with left PaCiG and PAG. FM patients also showed lower connectivity of the following seeds: right ACC with right SI; left AMYG with left CRBL; left AnG with left INS, right INS, right supramarginal gyrus (SMG), and left middle frontal gyrus (MidFG); left GP with left FP and left PCN; left INS with left PCN; right SMA with right TP, right M1, and left TP **(Figure 3, Table III).** These results show a disrupted rs-FC of the e-PNN in FM patients, when compared with HC. In our study, covariates did not show any significant influence in the main contrasts thus, rs-FC alterations in FM patients seem to be independent of age, years with FM diagnosis, as well as anxiety and depression symptoms.

**Figure 3.**
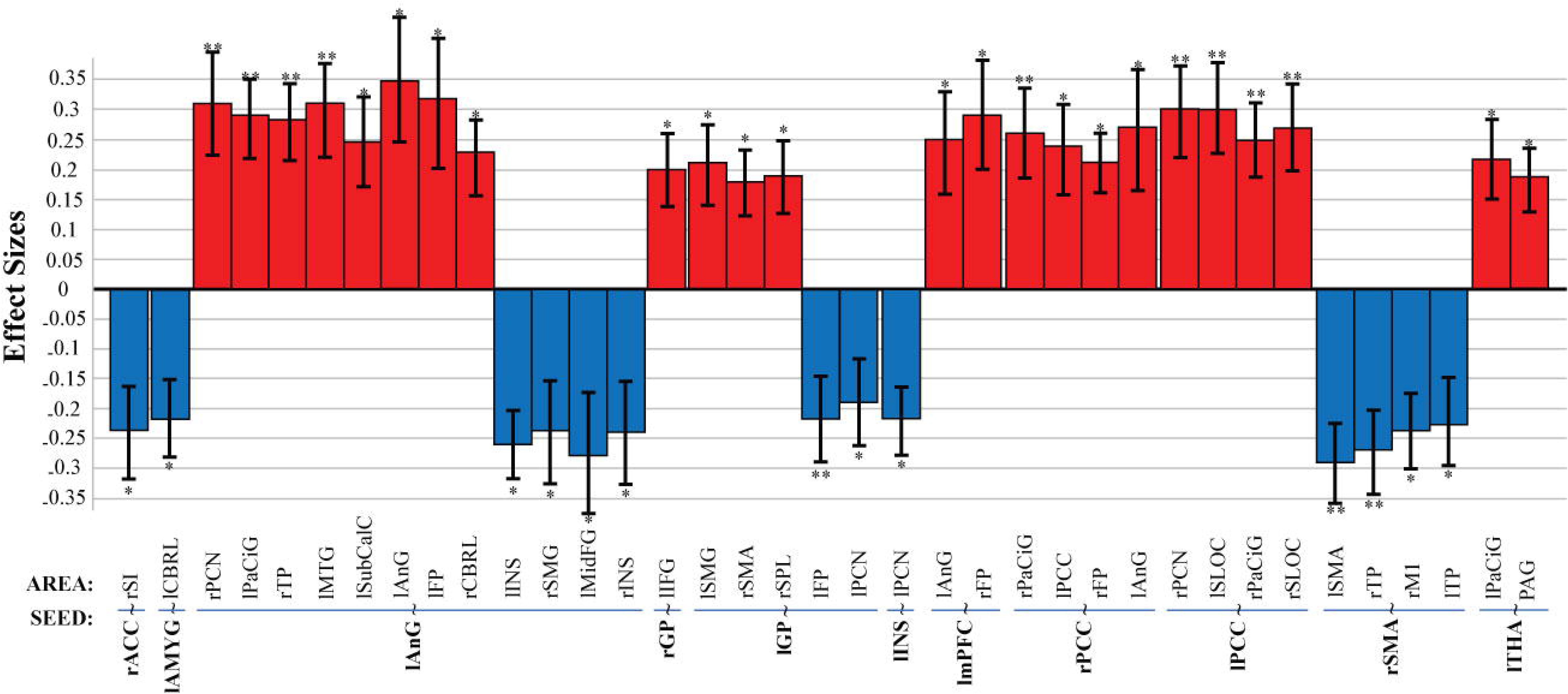
Baseline Condition. FM vs HC. Significant seed-to-voxel rs-fMRI functional connectivity of the e-PNN at rest; positive effects represent higher connectivity; negative effects represent lower connectivity. e-PNN, experimental pain neural network; FM, fibromyalgia; HC, healthy controls; l, left; r, right. Seeds: ACC, anterior cingulate cortex; AMYG, amygdala; AnG, angular gyrus; GP, globus pallidus; GP, globus pallidus; INS, insular cortex; mPFC, medial pre-frontal cortex; PCC, posterior cingulate cortex; PCC, posterior cingulate cortex; SMA, supplementary motor area; THA, thalamus. Correlated areas: AnG, angular gyrus; CRBL, cerebellum; FP, frontal pole; IFG, inferior frontal gyrus; INS, insular cortex; SLOC, superior lateral occipital cortex; MTG, medial temporal gyrus; MidFG, middle frontal gyrus; PaCiG, paracingular gyrus; PAG, periaqueductal gray matter; M1, precentral gyrus (primary motor cortex); SI, postcentral gyrus (primary somatosensory cortex); PCN, precuneus; PCC, posterior cingulate cortex; SubCalC, subcallosal cortex; SMG, supramarginal gyrus; SPL, superior parietal lobe; SMA, supplementary motor area; TP, temporal pole. *p<0.05, **p<0.001. All analyzed contrasts where corrected by multiple comparisons using the false discovery rate (FDR) at 0.05.

**Table III.**
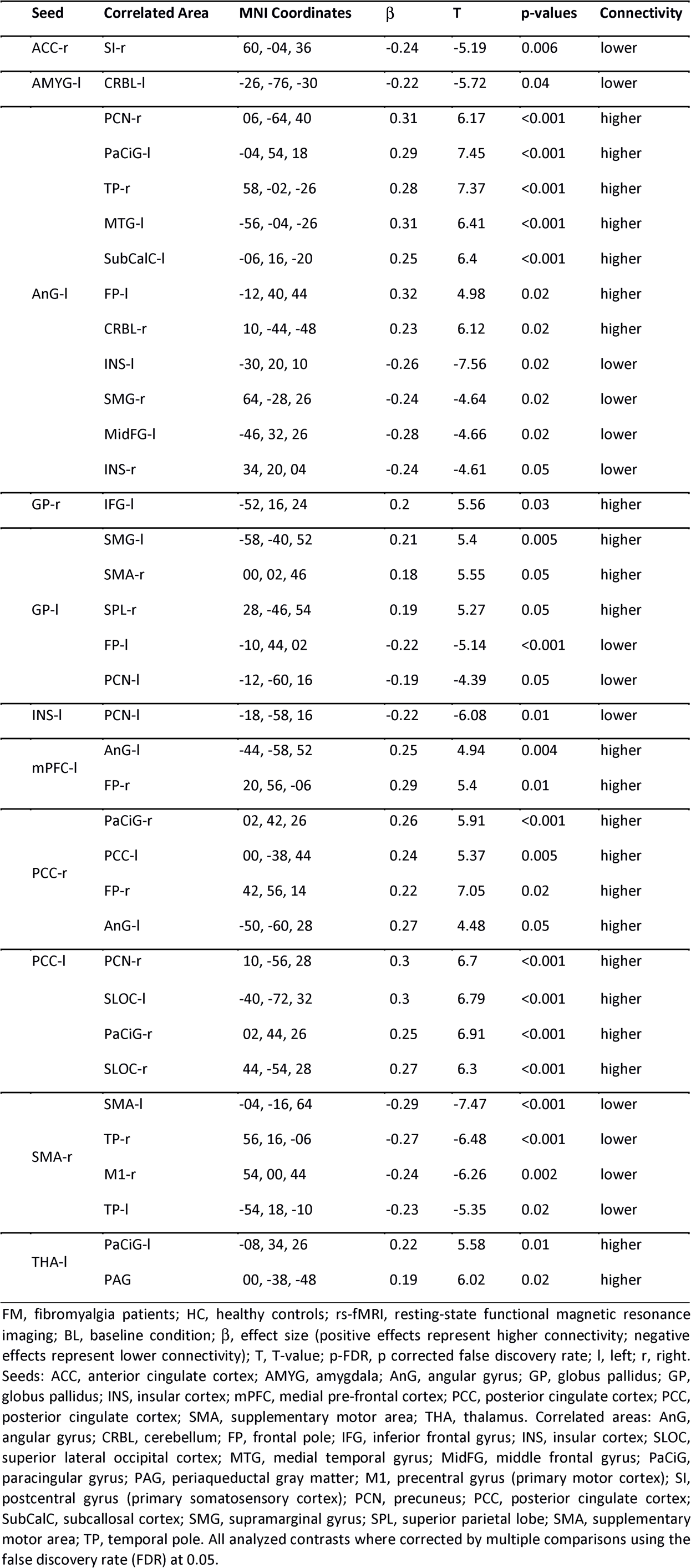
**Results of the paired t-tests of the FM vs HC rs-fMRI contrast analysis in the BL condition**. FM, fibromyalgia patients; HC, healthy controls; rs-fMRI, resting-state functional magnetic resonance imaging; BL, baseline condition; β, effect size (positive effects represent higher connectivity; negative effects represent lower connectivity); T, T-value; p-FDR, p corrected false discovery rate; l, left; r, right. Seeds: ACC, anterior cingulate cortex; AMYG, amygdala; AnG, angular gyrus; GP, globus pallidus; GP, globus pallidus; INS, insular cortex; mPFC, medial pre-frontal cortex; PCC, posterior cingulate cortex; PCC, posterior cingulate cortex; SMA, supplementary motor area; THA, thalamus. Correlated areas: AnG, angular gyrus; CRBL, cerebellum; FP, frontal pole; IFG, inferior frontal gyrus; INS, insular cortex; SLOC, superior lateral occipital cortex; MTG, medial temporal gyrus; MidFG, middle frontal gyrus; PaCiG, paracingular gyrus; PAG, periaqueductal gray matter; M1, precentral gyrus (primary motor cortex); SI, postcentral gyrus (primary somatosensory cortex); PCN, precuneus; PCC, posterior cingulate cortex; SubCalC, subcallosal cortex; SMG, supramarginal gyrus; SPL, superior parietal lobe; SMA, supplementary motor area; TP, temporal pole. All analyzed contrasts where corrected by multiple comparisons using the false discovery rate (FDR) at 0.05.

#### Auditory Conditions in Fibromyalgia Patients

The Cpre vs Cpos contrast analysis for FM patients was not significant, thus the control condition behaved as expected. However, the Mpre vs Mpos contrast analysis revealed that after listening to music, FM patients showed decreased connectivity of the left ACC with right posterior superior temporal gyrus (STG) and right SPL; the left AnG with right PCN, left superior frontal gyrus (SFG), right SFG, right PCC, and right posterior MTG; left INS with left M1; left M1 with left PaCiG; left SI with right occipital pole (OP); FM patients showed connectivity increase only of the left AMYG with right MidFG **(Figure 4a, Table IV).**

**Table IV.** **Results of the paired t-tests of the Mpre vs Mpos rs-fMRI contrast analysis in FM patients**. FM, fibromyalgia patients; Mpre, pre-music; Mpos, post-music; rs-fMRI, resting-state functional magnetic resonance imaging; β, effect size (positive effects represent increased connectivity; negative effects represent decreased connectivity; after listening to music); T, T-value; p-FDR, p corrected false discovery rate; l, left; r, right. Seeds: ACC, anterior cingulate cortex; AMYG, amygdala; AnG, angular gyrus; INS, insular cortex; M1, primary motor cortex; SI, primary somatosensory cortex. Correlated areas: MidFG, middle frontal gyrus; MTG, middle temporal gyrus; OP, occipital pole; PaCiG, paracingulate gyrus; M1, precentral gyrus (primary motor cortex); PCN, precuneus; PCC, posterior cingulate cortex; SFG, superior frontal gyrus; STG superior temporal gyrus; SPL superior parietal lobe. All analyzed contrasts where corrected by multiple comparisons using the false discovery rate (FDR) at 0.05.

| Seed | Correlated Area | MNI Coordinates | $\beta$ | T | p-values | Connectivity |
| --- | --- | --- | --- | --- | --- | --- |
| ACC-l | STG-r | 68, -16, 04 | -0.22 | -7.09 | <0.001 | Decreased |
|  | SPL-r | 18, -52, 72 | -0.21 | -6.22 | 0.04 | Decreased |
| AMYG-l | MidFG-r | 28, 30, 36 | 0.18 | 5.97 | 0.02 | Increased |
| AnG-l | SFG-r | 10, 16, 64 | -0.22 | -6.22 | 0.01 | Decreased |
|  | SFG-l | -08, 18, 60 | -0.22 | -7.64 | 0.03 | Decreased |
|  | PCC-r | 02, -16, 48 | -0.21 | -7.05 | 0.05 | Decreased |
|  | MTG-r | 70, -16, -08 | -0.25 | -7.97 | 0.05 | Decreased |
|  | PCN-r | 10, -56, 28 | -0.21 | -6.99 | 0.05 | Decreased |
| INS-l | M1-l | -18, -26, 56 | -0.18 | -8.72 | 0.03 | Decreased |
| M1-l | PaCiG-l | -04, 36, 26 | -0.2 | -6.86 | 0.02 | Decreased |
| SI-l | OP-r | 18, -92, 06 | -0.2 | -5.99 | 0.001 | Decreased |

FM, fibromyalgia patients; Mpre, pre-music; Mpos, post-music; rs-fMRI, resting-state functional magnetic resonance imaging; $\beta$ , effect size (positive effects represent connectivity increase; negative effects represent connectivity decrease; after listening to music); T, T-value; p-FDR, p corrected false discovery rate; l, left; r, right. Seeds: ACC, anterior cingulate cortex; AMYG, amygdala; AnG, angular gyrus; INS, insular cortex; M1, primary motor cortex; SI, primary somatosensory cortex. Correlated areas: MidFG, middle frontal gyrus; MTG,

#### Auditory Conditions in Healthy Controls

The Cpre vs Cpos contrast for HC was not significant for HC either. However, the Mpre vs Mpos contrast analysis, revealed that after listening to music HC showed increased connectivity of the right AMYG with right SI, left SLOC, and right SLOC; right AnG with left lingual gyrus (LG); decreased connectivity of the right INS with left CRBL; left PAG with left M1 and left SPL; increased connectivity of the right SI with right superior LOC and right hippocampus (HIPP); and increased connectivity of the right SII with right M1 **(Figure 4b, Table V).** These results also evidence music-related changes in the rs-FC of the e-PNN in FM patients and healthy controls. However, the connectivity patterns are different between the two groups, suggesting divergent effects on the e-PNN areas from listening to music not related to pain.

**Figure 4.**
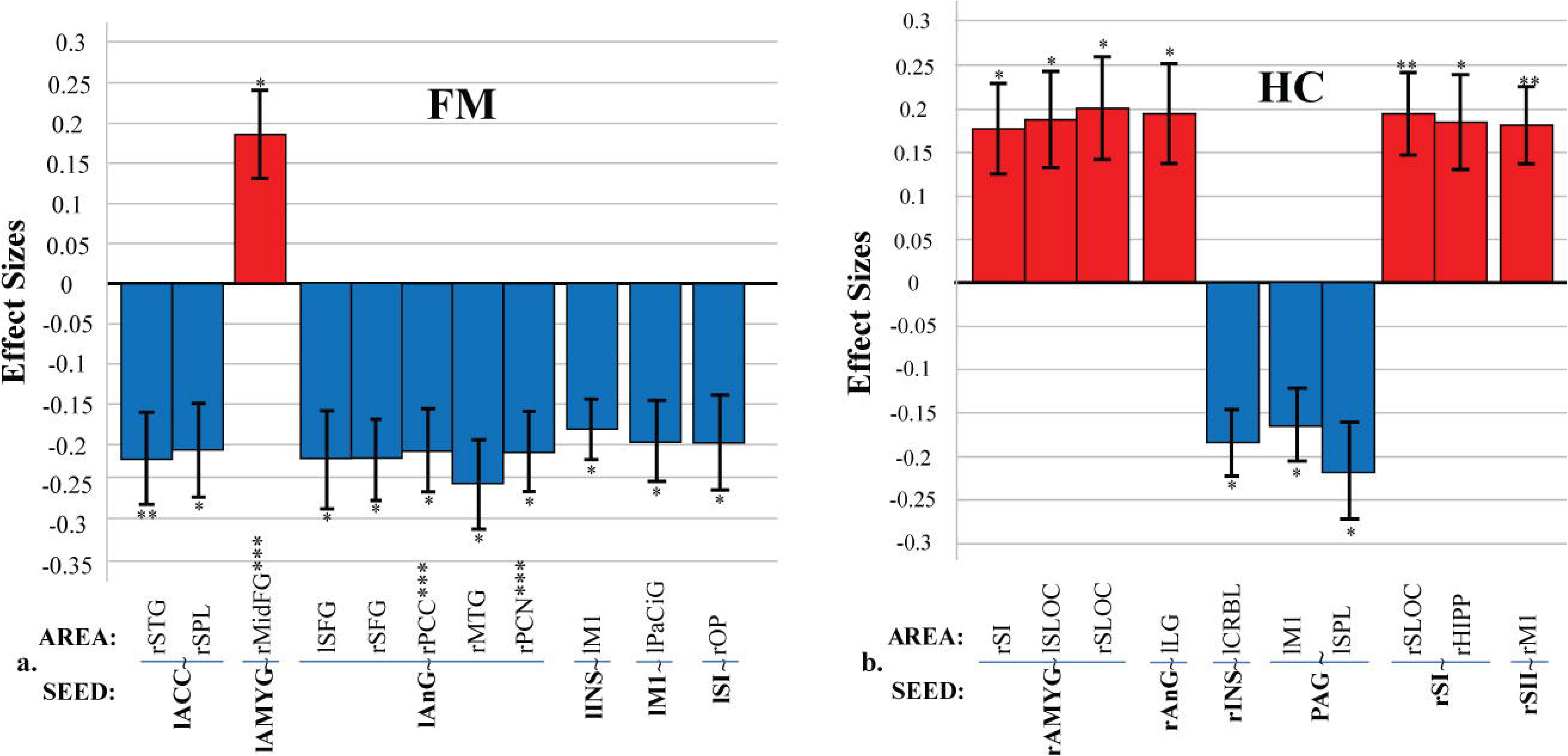
Mpre vs Mpos Contrast. Significant seed-to-voxel rs-fMRI FC of the e-PNN after listening to music; positive effects represent increased connectivity; negative effects represent decreased connectivity. **a. FM; b. HC.** e-PNN, experimental pain neural network; FM, fibromyalgia; HC, healthy controls; rs-fMRI, resting state functional magnetic resonance imaging; l, left; r, right. Seeds: ACC, anterior cingulate cortex; AMYG, amygdala; AnG, angular gyrus; INS, insular cortex; M1, primary motor cortex; PAG, periaqueductal gray matter; SI, primary somatosensory cortex; SII, secondary somatosensory cortex. Correlated areas: CRBL, cerebellum; HIPP, hippocampus; LG, lingual gyrus; MidFG, middle frontal gyrus; MTG, middle temporal gyrus; OP, occipital pole; PaCiG, paracingulate gyrus; M1, precentral gyrus (primary motor cortex); PCN, precuneus; PCC, posterior cingulate cortex; SI, postcentral gyrus (primary somatosensory cortex); SFG, superior frontal gyrus; SLOC, superior lateral occipital cortex; STG superior temporal gyrus; SPL superior parietal lobe. *p < 0.05, **p < 0.001, ***FC change correlated with pain scores (only ΔPI). All analyzed contrasts where corrected by multiple comparisons using the false discovery rate (FDR) at 0.05.

**Table V.**
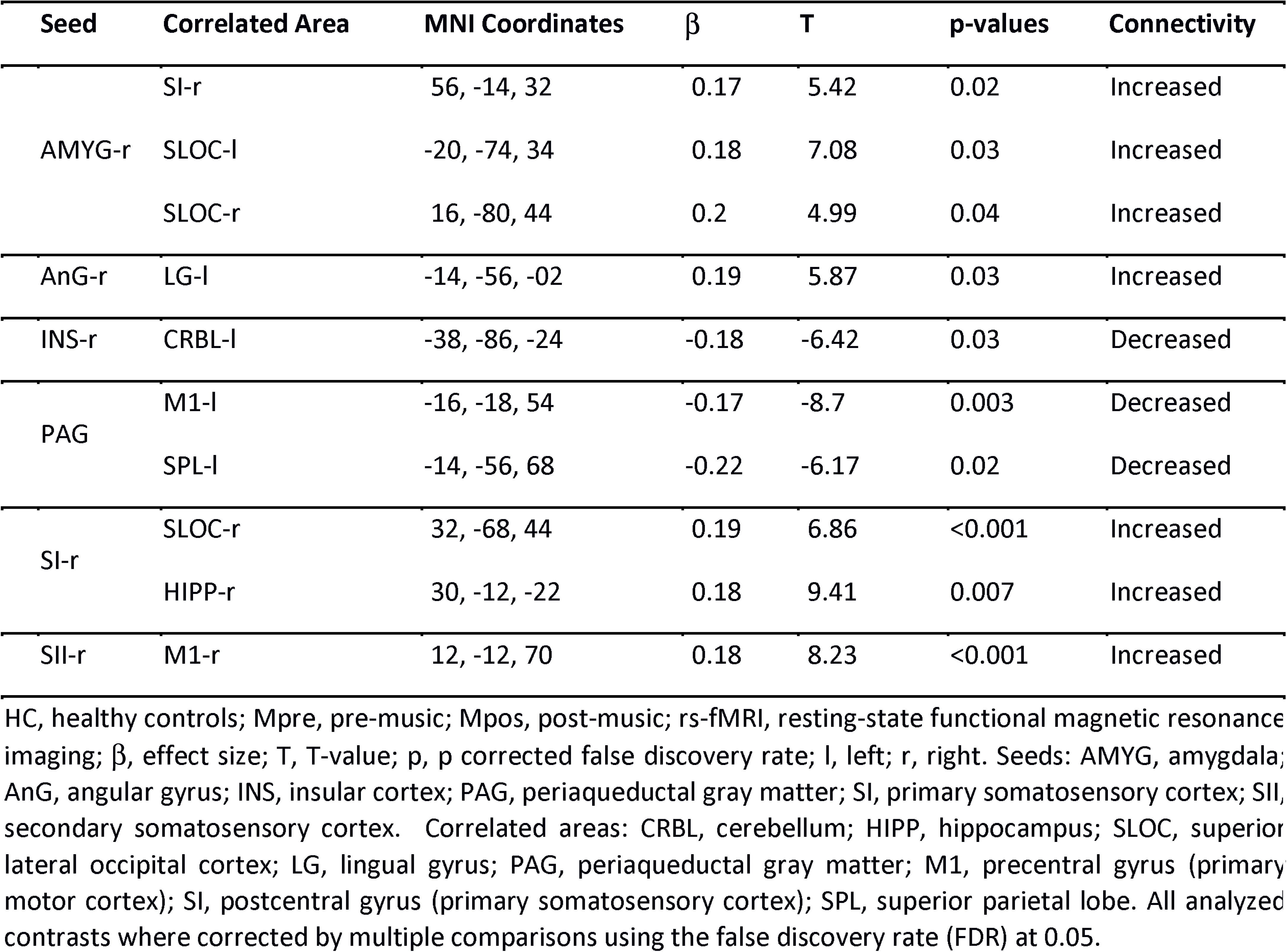
**Results of the paired *t*-tests of Mpre vs Mpos rs-fMRI contrast analysis in HC**. HC, healthy controls; Mpre, pre-music; Mpos, post-music; rs-fMRI, resting-state functional magnetic resonance imaging; β, effect size (positive effects represent increased connectivity; negative effects represent decreased connectivity; after listening to music); T, T-value; p, p corrected false discovery rate; l, left; r, right. Seeds: AMYG, amygdala; AnG, angular gyrus; INS, insular cortex; PAG, periaqueductal gray matter; SI, primary somatosensory cortex; SII, secondary somatosensory cortex. Correlated areas: CRBL, cerebellum; HIPP, hippocampus; SLOC, superior lateral occipital cortex; LG, lingual gyrus; PAG, periaqueductal gray matter; M1, precentral gyrus (primary motor cortex); SI, postcentral gyrus (primary somatosensory cortex); SPL, superior parietal lobe. All analyzed contrasts where corrected by multiple comparisons using the false discovery rate (FDR) at 0.05.

#### Neural Correlates of Music-Induced Analgesia in Fibromyalgia Patients

Significant correlations were found between the change of pain scores (ΔPI and ΔPU) and the change of rs-fMRI data for FM patients in the Mpre vs Mpos contrast (Δrs-FC). ΔPI was negatively correlated with the decrease of rs-FC between left AnG and right PCC (r = −0.28, p = 0.04), negatively correlated with the decrease of rs-FC between left AnG and right PCN (r = −0.49, p = 0.04), and positively correlated with the increase of rs-FC between left AMYG and right MidFG (r = 0.56, p = 0.02), after listening to music **(Figure 5).** In other words, the greater the analgesic effect, the greater the decrease of rs-FC between AnG, PCC and PCN, and the greater the increase of rs-FC between AMYG and MidFG. ΔPU did not show any significant correlation with rs-FC of the e-PNN after listening to music **(Supplementary Figure 2).**

**Figure 5.**
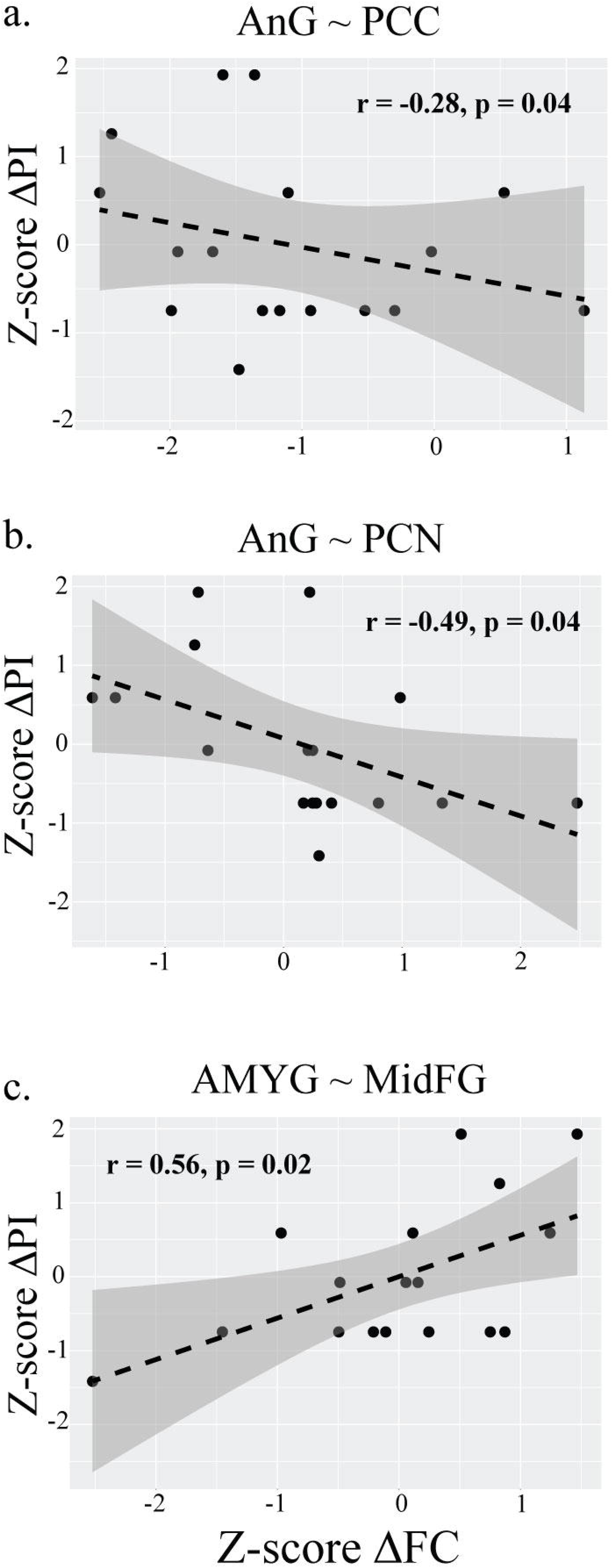
Scatterplot and regression line. Correlation between Mpre vs Mpos contrast (ΔFC) and ΔPI in FM patients, **a.** AnG˜PCC; **b.** AnG˜PCN; **c.** AMYG˜MidFG; FM, fibromyalgia patients; Mpre, before music; Mpos, after music; ΔPI, difference of pain intensity; ΔFC, difference in Mpre vs Mpos FC contrast; AMYG, amygdala; AnG, angular gyrus; PCC, posterior cingulate cortex; PCN, precuneus; MidFG, middle frontal gyrus; r, Pearson’s correlation coefficient.

## DISCUSSION

In our study, we investigated the neural correlates of music-induced analgesia in fibromyalgia patients, using rs-fMRI. We found that FM patients exhibited greater anxiety and depression symptoms, catastrophizing and perception of pain. Seed-based connectivity analysis showed a baseline disrupted resting state functional connectivity (rs-FC) of the pain neural network in FM patients. We also found that listening to music reduced pain in FM and that this analgesic effect was related to a decrease of the rs-FC of the left AnG with right PCC and right PCN, as well as an rs-FC increase of the left AMYG with right MidFG. FM patients show a myriad of psychiatric comorbidities including depression, anxiety, obsessive-compulsive disorder, and post-traumatic stress disorder, likely because there are common triggers (eg., early life stress or trauma), as well as shared pathophysiology (Uçar et al., 2015; Garcia-Fontanals et al., 2017; Costa et al., 2017). In addition, they experience a reduced self-regulatory capacity (Rost et al., 2017), leading to high scores on self-perception scales. Central sensitization of pain perception, in addition to a focused attention on their own pathological condition, may boost chronic pain in FM.

### Baseline Resting State Functional Connectivity Analysis

Our results showed baseline rs-FC differences in our experimental pain neural network (e-PNN) between groups. We showed that FM patients display significantly higher rs-FC of the AnG, PCC, GP, mPFC, and THA, compared to HC. The higher rs-FC of the AnG, PCC and mPFC with areas such as the PCN and SLOC, may evidence a dynamic coupling of the DMN’s rs-FC during pain perception, possibly secondary to a focused attention on their condition (Kucyi et al., 2017). FM patients may continuously engage autobiographical and self-awareness processes, commonly produced during rumination (Kucyi et al., 2014). Our results seem to be consistent with previous neuroimaging studies demonstrating that these regions are activated during pain perception (Apkarian et al., 2005) and affected in FM (Napadow et al., 2012). The higher rs-FC between the THA and the PAG may relate to the neuronal facilitation of pain input into the central nervous system (Staud, 2012; Truini et al., 2016; Potvin & Marchand, 2016). Previous studies have found several brain regions such as the PAG, INS, FP, AMYG, hypothalamus, and RVM, are involved in the descending pain modulatory system (DPMS) (Tracey et al., 2007; Henderson & Keay, 2017; Chen et al., 2017; Bannister & Dickenson, 2017). Thus, a higher rs-FC between the brain stem and the THA may support the hypothesized facilitation of pain perception, though functional connectivity does not convey directionality or the type of neuronal function involved (excitation or inhibition) (Rogers et al., 2007; Vattikonda et al., 2016). The higher rs-FC of basal ganglia (GP) with SMG and SMA may play a key role in the integration of motor, emotional, autonomic and cognitive aspects of pain in FM, with an enhanced function of areas related to pain processing (Cifre et al., 2012).

Additionally, we found lower rs-FC of the ACC, AMYG, AnG, INS and SMA in FM patients. The lower rs-FC of the right SMA with M1, contralateral SMA and bilateral TP may explain a disrupted connectivity of motor areas with limbic and paralimbic regions. A proposed explanation for this is that the TP binds complex, highly processed perceptual inputs to visceral emotional responses (Olson et al., 2007). FM patients seem to process emotion and pain in a different manner than the general population (Montoya et al., 2005; Mhalla et al., 2010), with relevant deficits in affective modulation measured by cardiac responses, heart rate variability, and neuroimaging (Rosselló et al., 2015), suggesting alteration of emotional and attentional aspects of information processing in chronic pain (Sitges et al., 2007). Moreover, lower rs-FC between other emotion related regions such as AMYG and INS may support this hypothesis (Ichesco et al., 2015; Lazaridou et al., 2017). Finally, the lower rs-FC of ACC and AnG with INS and SI may relate to an altered somatosensory processing with limbic and pain related areas in FM patients (Kim et al., 2014; Loggia et al., 2015; Schreiber et al., 2017), which might be characterized by dissociation between sensory and affective components of pain-related information (Sitges et al., 2007). Somatic dysfunction in FM, including clinical pain, pain catastrophizing, autonomic dysfunction, and temporal summation, are closely related with the degree to which pain alters SI connectivity with affective pain processing regions (Kim et al., 2015), and suggests that affective mood states can modulate central excitability thresholds in chronic pain states (Montoya et al., 2005). Overall, it seems that FM patients feature patterns of connectivity in subcortical and cortical pain related areas that are mostly consistent across studies and that evidently reflects their physiopathology.

### Music Effects on Pain and Functional Connectivity

We found that listening to music reduced pain in FM patients. We also found significant changes in rs-FC after listening to music and not after noise in both groups. This suggests that our control condition worked as expected (no effect) and rs-FC changes in the music condition cannot be attributed solely to sound perception. Although the seeds were selected based on prior evidence related to pain perception (Gracely et al., 2002; Zaki et al., 2007; Baliki et al., 2008; Burgmer et al., 2009), these regions are not exclusively related to pain, i.e. insula (Xue et al., 2010; Jakab et al., 2012). In FM after listening to music, there was a left lateralized decreased rs-FC mainly in ACC, AnG, INS, M1 and SI connectivity, and an increase in AMYG. The ACC has been described as a main hub in cognitive control, from reward processing and performance monitoring, to the execution of control and action selection (Shenhav et al., 2012) and even suggested to be specific to pain processing (Lieberman & Eisenberger, 2015). There is evidence that the ACC is involved in processing the affective and unpleasant aspects of pain (Gracely et al., 2002). The decreased rs-FC of the ACC with the STG (primary auditory cortex) after listening to music suggests an influence of music and sound processing in the modulation of pain that may not be explained solely by distraction, as the rs-fMRI was acquired after the music listening (Harriott & Schwedt, 2014; Schwedt et al., 2015). Music-evoked memories may play a role in the sustained distraction that prevails with the analgesic effect (Koelsch, 2015). Similarly, the decreased rs-FC of ACC with SPL (somatosensory association) suggests disentanglement between these areas closely related to pain at the cortical level that may be related to the analgesic effect (Orenius et al., 2017). The decrease of rs-FC between INS-M1, and M1-PaCiG, strongly suggests an analgesic effect, as these regions are usually activated during pain perception (Orenius et al., 2017). The decrease of rs-FC between AnG and SFG (premotor and SMA) may suggest disengagement of areas related to pain and attention, after listening to music (Seghier, 2013; Greicius et al., 2003). The decrease of FC between AnG, PCC and PCN, may suggest a decrease in the activity of the DMN after listening to music, which may be related to the central analgesic effect in FM. The AnG has been related to several brain functions including semantic processing, word reading and comprehension, number processing, memory retrieval, attention and spatial cognition, reasoning, and social cognitions (Andrews-Hanna, 2012; Seghier, 2013). Additionally, it has been shown to be an important hub of the DMN, connecting perception, attention and spatial cognition during mental navigation at rest (Fox et al., 2005). Therefore, the decreased rs-FC may suggest a temporary disruption of autobiographical memory and self-awareness of the pain. Finally, the increase of rs-FC between AMYG and MidFG after music listening may be secondary to the association between auditory attention (Nakai et al., 2005), memory retrieval (Ranganath et al., 2003), and positive emotions (Kerestes et al., 2012), consistent with the use of a known, pleasant and emotionally positive music track. It is important to mention that FM patients show an altered baseline rs-FC (Cifre et al., 2012) that seems to be the result of the chronic pain and/or disease. Consequently, the dysfunction in the DPMS may induce a reorganization of the "pain-analgesia network", and in order to modulate pain and produce an analgesic effect the system may be utilizing other circuits rather than straightforward ones in FM.

Listening to music also showed significant changes in rs-FC of the e-PNN in healthy controls (Karmonik et al., 2016; Brodal et al., 2017; Alluri et al., 2017). However, the connectivity patterns show solely the after effects of listening to music and differ from those identified in the FM patients, starting with a right lateralization of the significant changes, compared to the left lateralization in FM. Our results showed both increase and decrease rs-FC of selected seeds for the e-PNN after listening to music, such as AMYG, AnG, INS, PAG, SI, and SII. As mentioned in the beginning of this discussion, the areas selected to build the e-PNN are not exclusive for pain processing, and are active during other cognitive processes. The emotional valence of music may play a role in connecting areas related to limbic, somatomotor, memory and visual imagery processes (Koelsch, 2014). The intensity of pleasure experienced from music listening suggests a relation with dopamine reward system of the brain, and neural activity in surrounding limbic regions, indicative of emotional arousal (Salimpoor et al., 2009; Kringelbach, 2005; Kringelbach & Berridge, 2010). AMYG showed an increased rs-FC with SI and SLOC in both hemispheres after listening to music, parietal areas of somatosensory functions and occipital areas involved in visual mental imagery (Platel et al., 1997). We found increased rs-FC of the AnG with the LG, possibly secondary to visual memory and visuo-limbic processes engaged after listening to music (Rogenmoser et al., 2016). The rs-FC increase of somatosensory cortices (SI and SII) and occipital association cortices with motor cortex and hippocampus support this hypothesis (Groussard et al., 2014; Frühholz et al., 2016). We also found decrease of rs-FC between INS and PAG with motor areas such as M1 (primary motor cortex), CRBL, and SPL, which may be caused by a focused attention on music, and a probable state of relaxation in healthy adults (Brattico et al., 2017).

### Limitations

The FM patients in this study were under different types of medication and had several comorbidities that we could not control, and which may affect baseline rs-FC. However, our main results show an analgesic effect of self-chosen, pleasant familiar music, related to changes in rs-FC. To account for any possible comorbidity confounds, we analyzed the data using depression and anxiety covariates, where we found no significant effect of such (see Supplementary Methods and Results). It should be noted that the dosage of music intervention in this study may be limited (5 min), and the precise duration of the analgesic effect may be variable; though it seems 5 minutes was enough to elicit MIA. Our FM patients were not blinded to the music condition, which can produce a bias. Given the nature of music, blinding for participants is near impossible and thus a control such as white noise is used instead. In fact, our control condition (pink noise) behaved as expected, as we did not find any significant effect of it on pain perception or rs-FC analyses. Finally, it should be noted that our HC group did not experience pain, thus, a comparative measure to pain was not possible.

## CONCLUSIONS

Our results show that FM patients experience an analgesic effect after listening to music, and this effect correlates with changes in resting state functional connectivity (rs-FC) of our chosen pain network. Specifically, the analgesic effect (greater ΔPI) was correlated with decreased rs-FC of lAnG with rPCC and rPCN, and with increased rs-FC of lAMYG with rMidFG. Hence, music-induced analgesia correlated with rs-FC changes between important areas of the DMN regions processing emotion, memory retrieval, and auditory attention. We therefore suggest cognitive and emotional modulation of pain in FM after listening to music. MIA may arise as a consequence of top-down modulation, probably originated by distraction, relaxation, positive emotion, or a combination of these mechanisms.

## ACKNOWLEDGEMENTS

We would like to thank the group Fibromyalgia Queretaro for their help with our study. Thanks to Luis Concha-Loyola for the help with the MRI sequences and the methodological and technical support and Juan J. Ortiz for technical support. Thanks to Juan I. Romero-Romo for the help with the clinical logistics and with the recruitment of the patients. We would also like to thank Micah Allen for his valuable insight and suggestions about the experimental design. The authors declare no conflict of interest. This study was funded by the Funding for Neurology Research, the Augustinus Fonden, the Danish Basic Research Foundation, the Ulla and Mogens Folmer Andersden Foundation, and the Institute of Neurobiology UNAM.

---

**Supplementary Figure 1.** Connectivity maps of the e-PNN for FM (left) and HC (right) respectively. Colors show either positive (red–yellow) or negative (blue–light blue) correlations of the e-PNN (mean of all seeds) with the Default Mode Network (DMN), with areas such as mPFC, PCC, and AnG, among others.

**Supplementary Figure 2. Scatterplot and regression line. Correlation between Mpre vs Mpos contrast (ΔFC) and ΔPU in FM patients.** FM, fibromyalgia patients; Mpre, before music; Mpos, after music; ΔPU, difference of pain unpleasantness; ΔFC, difference in Mpre vs Mpos FC contrast; AMYG, amygdala; AnG, angular gyrus; PCC, posterior cingulate cortex; PCN, precuneus; MidFG, middle frontal gyrus; r, Pearson’s correlation coefficient.

**Supplementary Table I.** MNI Coordinates of Seeds’ Center for the e-PNN. MNI, Montreal Neurological Institute; e-PNN, experimental pain neural network; ACC, anterior cingulate cortex; AMYG, amygdala; AnG, angular gyrus; BA41, primary auditory cortex; CAU, caudate; GP, globus pallidus; INS, insular cortex; M1, primary motor cortex; mPFC, medial prefrontal cortex; PAG, periaqueductal gray matter; PCC, posterior cingulate cortex; PUT, putamen; SI, primary somatosensory cortex; SII, secondary somatosensory cortex; SMA, supplementary motor area; STS, superior temporal sulcus; THA, thalamus. a, Gracely et al., 2002; b, Harvard-Oxford atlas FSLview; c, Juelich Histological atlas FSLview; d, Baliki et al., 2008; e, Zaki et al., 2007; f, Burgmer et al., 2009.

